# LC3 and STRAP regulate actin filament assembly by JMY during autophagosome formation

**DOI:** 10.1101/451393

**Authors:** Xiaohua Hu, R. Dyche Mullins

**Affiliations:** Department of Cellular and Molecular Pharmacology UCSF School of Medicine Howard Hughes Medical Institute

## Abstract

During autophagy actin filament networks move and remodel cellular membranes to form autophagosomes that enclose and metabolize cytoplasmic contents. Two actin regulators, WHAMM and JMY, participate in autophagosome formation, but the signals linking autophagy to actin assembly are poorly understood. We show that, in non-starved cells, cytoplasmic JMY co-localizes with STRAP, a regulator of JMY’s nuclear functions, on non-motile vesicles with no associated actin networks. Upon starvation, JMY shifts to motile, LC3-containing membranes that move on actin comet tails. LC3 enhances JMY’s *de novo* actin nucleation activity via a cryptic actin-binding sequence near JMY’s N-terminus, and STRAP inhibits JMY’s ability to nucleate actin and activate the Arp2/3 complex. Cytoplasmic STRAP negatively regulates autophagy. Finally, we use purified proteins to reconstitute LC3‐ and JMY-dependent actin network formation on membranes, and inhibition of network formation by STRAP. We conclude that LC3 and STRAP regulate JMY’s actin assembly activities *in trans* during autophagy.

**eTOC Blurb:** The actin regulator JMY creates filament networks that move membranes during autophagy. We find that, in unstarved cells, JMY is inhibited by interaction with the STRAP protein, but upon starvation JMY is recruited away from STRAP and activated by LC3.

## Introduction

Autophagy is a catabolic process during which membranes move and remodel to form autophagosomes: organelles bounded by double membranes that engulf and metabolize cytoplasmic contents (Shibutani and Yoshimori, 2014). In 2015 Mi et al. found evidence that branched actin networks created by the Arp2/3 complex drive some of the membrane movements required for autophagosome formation (Mi et al., 2015). Following this initial discovery, two groups identified a pair of related nucleation-promoting factors, WHAMM (WASP Homolog Associated with Membranes and Microtubules) and JMY (Junction Mediating and regulatorY protein), as contributors to autophagosome biogenesis. Kast et al. found that WHAMM promotes assembly of actin networks on early autophagosomal membranes and that decreasing WHAMM expression reduces the number and size of autophagosomes (Kast et al., 2015). Coutts and La Thangue found that JMY promotes autophagy by directing actin assembly on membranes that also contain the autophagy regulator, LC3. These authors also noticed that the N-terminal region of JMY contains a consensus LC3-interacting region (LIR) found in many autophagy-related proteins (Coutts and La Thangue, 2015). Intriguingly, removal of the N-terminal LIR not only disrupted the association of JMY with LC3-containing membranes, but also impaired JMY’s ability to create actin structures in the cytoplasm. The effect of N-terminal deletions on actin assembly is surprising, given that JMY’s previously described actin‐ and Arp2/3-binding sites are all located near the C-terminus. Little is known about how upstream factors regulate WHAMM‐ or JMY-directed actin assembly, but the results of Coutts and Lathangue (2015) suggest that JMY’s N-terminal region might regulate both localization and nucleation activity.

JMY is an enigmatic actin regulator. Initially described as a co-activator of p53-mediated apoptosis following DNA damage (Shikama et al., 1999), we later found that JMY contains an Arp2/3-activating sequence, called a WCA domain, common to Class II nucleation promoting factors (Zuchero et al., 2009; Welch and Mullins, 2002). This WCA domain resides in the C-terminal region of JMY and comprises three conserved motifs: (1) a set of three actin-binding Wasp-Homology 2 (WH2 or W) domains; (2) a ‘central connecting’ domain (C) that binds both actin and the Arp2/3 complex; and (3) an acidic region (A) that binds the Arp2/3 complex. In addition to promoting Arp2/3-dependent nucleation, JMY’s three WH2 domains also nucleate actin filaments on their own (Zuchero et al., 2009) using a mechanism similar to that of spire (Quinlan et al., 2005). With the possible exception of Las17p in budding yeast (Urbanek et al., 2013), this combination of intrinsic and Arp2/3-mediated nucleation activities is unique to JMY, and suggests that its function might be highly specialized.

Although it can function as a co-activator of p53-dependent transcription in the nucleus, JMY localizes predominantly to the cytoplasm in unperturbed cells (Firat-Karalar et al., 2011; Schlüter et al., 2014), where, in addition to autophagy, it has been reported to participate in a variety of cellular processes, including cell migration and adhesion (Coutts et al., 2009), regulation of neurite outgrowth (Firat-Karalar et al., 2011), and asymmetric cell division of mouse oocytes (Sun et al., 2011). JMY has also been reported to interact with endoplasmic reticulum-resident protein VAP-A (Schlüter et al., 2014), which may be important for autophagosome biogenesis since the endoplasmic reticulum (ER) is a major source of membranes for autophagosome formation (Shibutani and Yoshimori, 2014).

The regulation of JMY’s actin nucleation activity in the cytoplasm is less understood than the regulation of its nuclear functions. No modulators of JMY’s actin nucleation activity have been identified, while the acetyltransferase p300 and its binding partner STRAP (stress-responsive activator of p300, also named TTC5) are known to interact with JMY in the nucleus. STRAP appears to act as a scaffold, facilitating formation of the JMY/p300 complex and preventing JMY’s degradation by MDM2 (Demonacos et al., 2001). Little is known about the cytoplasmic role of STRAP, including whether it interacts with JMY in the cytoplasm.

In the present study we identify both positive and negative regulators of JMY’s actin nucleation activity in the cytoplasm. We show that LC3 directly recruits JMY to membrane surfaces and enhances its *de novo* actin nucleation activity via a cryptic actin-binding sequence near JMY’s N-terminus. Because the nucleation activity of the Arp2/3 complex requires pre-existing, ‘mother’ filaments, the enhancement of JMY’s N-terminal, cryptic nucleation activity has a knock-on effect that dramatically enhances actin network formation. We also show that STRAP potently inhibits actin nucleation stimulated by JMY and antagonizes JMY activation by membrane-associated LC3. STRAP also turns out to be a previously unrecognized negative regulator of starvation-induced autophagy. In fed cells JMY primarily associates with non-motile, STRAP-containing vesicles. Upon starvation-induced autophagy, JMY localization shifts to LC3-containing membranes which move on polarized actin networks.

## Results

### JMY translocates to motile LC3-positive vesicles from non-motile STRAP vesicles upon starvation-induced autophagy

To better understand the regulation of JMY activity in the cytoplasm we co-expressed a JMY-mCherry fusion protein together with two JMY interacting proteins, GFP-LC3B and SNAP-tagged STRAP in U2OS cells (Movie 1). In normally fed cells, both JMY and STRAP localized primarily to cytoplasmic foci (Figure 1A, 1E, S1A-C), and many JMY foci also contained STRAP (81% ± 18%, Figure S1A). We observed that only a small fraction of LC3 vesicles also contain JMY (6.9% ± 13%, Figure S1B). Upon induction of autophagy by starvation, more JMY puncta accumulated LC3 (from 10 ± 20% to 30 ± 24%, Figure S1A) while the overlap between JMY and STRAP sharply decreased (from 81 ± 18% to 34 ± 21%, Figure S1A), consistent with a shift in JMY’s location and function upon starvation (Figure 1A, 1E, movie 1). Moreover, vesicles containing JMY and LC3, but lacking STRAP began moving rapidly and persistently through the cytoplasm (0.29 ± 0.08 μm/sec) (Figure 1B, 1F). In contrast, vesicles containing JMY and STRAP but lacking LC3 were non-motile (0.003 ± 0.012 μm/sec) (Figure 1C, Kymograph 1, 1F).

**Figure 1.**
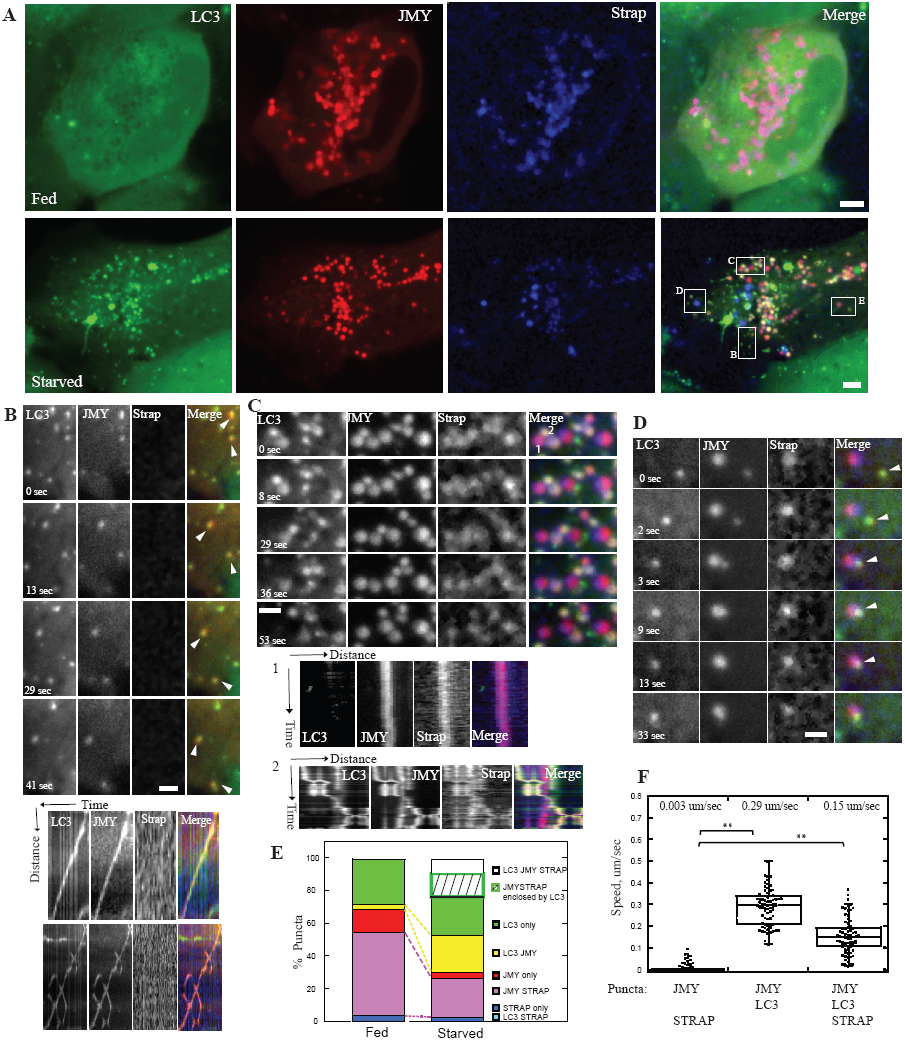
In U2OS cells JMY translocates from STRAP-positive non-motile foci to motile LC3-positive motile foci upon starvation. (A) JMY (red) and STRAP (blue) colocalize on foci under normally fed condition (top panel). More JMY (red) colocalizes with LC3 (green) when cells are starved in HBSS (bottom panel). (B) Two vesicles (arrowheads) that contain both JMY and LC3 but no STRAP (yellow color in merge channel) move persistently through cytoplasm. Kymographs show travel distance versus time. (C) Enlarged box indicates two populations of vesicles. 1).Vesicles that contain JMY and STRAP but no LC3 (magenta color in merge channel) are non-motile. Kymograph 1 shows one example of a non-motile JMY-and STRAP-positive vesicle. 2). Vesicles that contain all three proteins, JMY, STRAP and LC3 (yellow-white color in merge channel) move in a saltatory manner. Kymograph 2 shows one example of the saltatory movement of a JMY-, STRAP‐ and LC3-positive vesicle. (D) JMY‐ and LC3-positive vesicle (yellow color in merge channel) moves and fuses with JMY‐ and STRAP-(magenta color in merge channel) positive vesicle (arrowhead). The fused vesicle is non-motile during observation. (E) Quantification of colocalization of JMY, STRAP and LC3 under fed and starved condition. The percentage is normalized to total number of puncta (including all LC3, JMY and STRAP puncta) of the corresponding cell (n = 3 independent experiments, 24 cells, total number of puncta per cell: 15 ‐ 83). Quantifications of colocalization normalized to each protein are shown in Figure S1A-C. (F) Quantification of migration speed of JMY positive vesicles (Number of JMY-/LC3-positive puncta: 73; number of JMY-/STRAP‐ positive puncta: 208; number of JMY-/LC3-/STRAP‐ positive puncta: 73, n = 3, **p < 0.01, student t test). Temperature: 37⁰. Scale Bar: whole cell, 5 μm; zoom in box, 2 μm.

LC3 and STRAP almost never colocalized in the absence of JMY (from 0% in fed cells to 0.8 ± 2% in starved cells) (Figure S1B). Occasionally in starved cells, all three proteins, JMY, STRAP and LC3 were present on the same puncta (12 ± 9%, Figure S1A), and these puncta exhibited saltatory movement (0.15 ± 0.08 μm/sec) (Figure 1C, Kymograph 2, 1F), suggesting that LC3 and STRAP were competing to bind JMY and antagonistically modulating its activity. We also observed that some JMY and STRAP puncta (19% ± 17%, Figure S1A) were enclosed in LC3 vesicles (Figure S1D-E), and that these were much less motile than other JMY-LC3 vesicles (Figure S1F). In rare cases, we observed JMY‐ and LC3-positive vesicles migrate toward and merge with non-motile JMY‐ and STRAP-positive vesicles (Figure 1D), which might reflect a process in which LC3-containing membranes recruit JMY away from its interaction with STRAP.

We further confirmed the above result by co-expressing only two proteins at a time in U2OS cells: either GFP-LC3B and JMY-mCherry or JMY-mGFP and STRAP-mCherry. As in the experiments described above, JMY puncta displayed directed motility when colocalized with LC3, but showed little or no motility when co-localized with STRAP (Figure 2E, Figure S2, Movie 2, Movie 3). Taken together, our data indicate that starvation triggers JMY to shift from non-motile STRAP-containing membranes to motile, LC3‐ positive autophagosomes.

**Figure 2.**
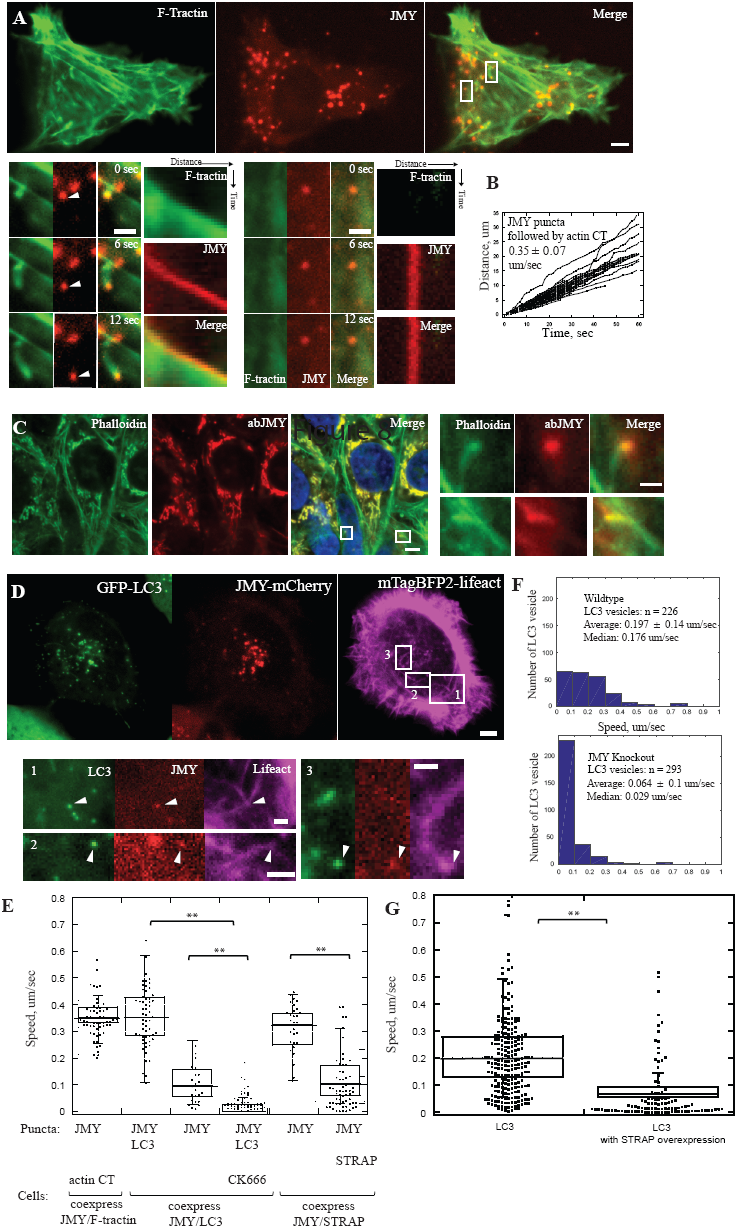
In U2OS cells LC3 and JMY vesicles move on polarized actin network. (A) A subset of punctate JMY structures (red) are propelled through the cytoplasm (*arrowheads*) by actin comet tails (green) in fed U2OS cells. Enlarged boxes show one motile JMY focus with an associated actin comet tail (left) and one non-motile JMY focus with no actin network (right). (B) Raw distance versus time plots of JMY-positive foci propelled by actin comet tails (63 puncta, 9 cells, n = 3 independent experiments). (C) Polarized actin networks (labeled by phalloidin, green) attach to endogenous JMY (JMY antibody, red) puncta in immunofluorescent staining in fed cells. (D) JMY/LC3-positive vesicles are propelled by polarized actin networks in cells starved in HBSS. Enlarged boxes show actin networks (magenta) pushing JMY (red) and LC3 (green) positive vesicles (arrowhead) toward plasma membrane (1), toward nucleus, (2) or in a more complex trajectory (“V” shape movement) (3). (E) Quantification of migration speed of JMY positive vesicles in cells expressing various sets of two proteins: JMY and F-tractin, JMY and LC3 (with or without CK666), JMY and STRAP (8∼11 cells, **p < 0.01, student t test) (F) Histograms show decrease in migration speed of LC3-positive vesicles in JMY knockout cell line (*bottom*) compared to wildtype U2OS cells (*top*). (G) The migration speed of LC3 vesicles decreases when U2OS cells over-express STRAP (20 ∼ 25 cells, n = 3 independent experiments, **p < 0.01, student t test). Temperature: 37⁰. Scale bar: whole cell, 5 μm; zoom in box, 2 μm.

### JMY promotes formation of actin networks on motile, LC3-positive vesicles

Coutts and Lathangue found actin associated with LC3‐ and JMY-positive vesicles, thus we hypothesized that JMY might assemble actin comet tails to drive vesicle motility (Coutts and La Thangue 2015). To test this possibility, we co-expressed JMY-mCherry with a live-cell probe for filamentous actin (GFP-F-tractin) in U2OS cells. Over half of JMY puncta (66 ± 16%, 293 puncta, from 9 cells) were associated with polarized actin comet tails, while the rest had no clear association with filamentous actin (Figure 2A). The actin-associated JMY foci moved rapidly through the cytoplasm, at an average speed of 0.35 ± 0.07 μm/sec (Figure 2A-B, 2E), apparently propelled by actin filament assembly (Movie 4) in a manner similar to the motility of some intracellular pathogens such as *Listeria monocytogenes*. To test whether these motile puncta were created by over-expression of labeled JMY we performed fluorescence microscopy on fixed U20S cells stained with phalloidin and an anti-JMY antibody to visualize the distribution of endogenous JMY and actin. As in the live cell experiments, we observed polarized actin networks associated with endogenous JMY foci (Figure 2C), indicating that JMY-associated actin comet tails are not an over-expression artifact.

To verify that movement of LC3/JMY foci is driven by polarized actin comet tails, we coexpressed fluorescent derivatives of JMY and LC3 together with a fluorescent probe for filamentous actin (mTagBFP-lifeact) in U2OS cells. Using three-color, fluorescence microscopy we found that all motile, JMY/LC3-positive vesicles are associated with actin comet tails (Figure 2D, movie 5). Slightly more than half of the motile LC3/JMY foci (56%) migrated toward the plasma membrane, while 21.9% migrated toward the nucleus, and the remainder moved in a more circumferential path around the cell. It is likely that these autophagosomes are movingalong ER tubule because, we observed JMY comigrate with its binding protein VAPA, an ER-resident protein involved in vesicle budding and membrane transport (Schlüter et al., 2014). In these experiments, polarized actin network propel JMY moving along VAP-A labeled ER tubule (Figure S3). When we treated cells with CK666,which inhibits nucleation activity of Arp2/3 complex, all of the JMY/LC3-positive puncta became non-motile, with an average speed of 0.015 ± 0.028 μm/sec (Figure 2E, Movie 6), indicating that movement of JMY/LC3 positive structures requires Arp2/3-mediated actin filament assembly.

To investigate JMYs role in promoting actin-driven movement of LC3-containing vesicles, we used CRISPR to edit the JMY gene locus and knockout JMY protein expression in our U2OS cells (Figure S4A-B). Upon starvation we observed that the majority of LC3-positive puncta (75.2 % ± 21%) in these JMY knockout cells remain non-motile (Figure 2F, Movie 7). We interpret this result as evidence that JMY-induced actin assembly drives motility of LC3-containing membranes.

We next treated wildtype U2OS cells with Bafilomycin A, to prevent acidification of mature autophagosomes and block autophagic flux (Klionsky et al., 2008; Yamamoto et al., 1998). Under these conditions, JMY/LC3-positive vesicles accumulated in the perinuclear region (Figure S4D). Interestingly, when we added the Arp2/3 inhibitor CK666 together with Bafilomycin A, JMY/LC3 positive vesicles lost their perinuclear localization (Figure S4E-F), suggesting that, in addition to microtubule motors (Pankiv and Johansen, 2010; Pankiv et al., 2010; Jahreiss et al., 2008), actin-based movement may also be important for bulk centripetal flow of autophagosomes and/or detachment from their original sites of biogenesis.

### Co-localization of JMY with other autophagosomal markers

To better define JMY’s role in autophagy we looked for co-localization with other membrane markers in normal and perturbed cells. An early step in autophagy is the phosphorylation of phosphatidylinositol (PI) to form PI(3)P, which then recruits LC3. We found that blocking the activity of PI-kinases with wortmannin A disrupted the punctate localization of both LC3 and JMY (Figure S5A), suggesting that JMY recruitment is not upstream of PI(3)P accumulation. We did, however, observe colocalization and co-migration of JMY with DFCP1 and ATG9, two proteins involved in the initiation and nucleation phases of phagophore assembly (Figure S5B-C). We also investigated late stages of autophagy, when autophagosomes fuse with lysosomes and their contents are degraded, and we found that JMY colocalizes with the late endosome and lysosome marker, Lamp1 (Figure S5D). Coutts and Lathangue (2016) previously reported that JMY does not co-localize with Lamp1, but we find the association most obvious when cells are treated with bafilomycin A to inhibit autolysosome maturation. These results indicate that JMY associates with autophagosomal membranes from nucleation to final fusion with lysosomal or endosomal membranes.

### The JMY-binding protein STRAP regulates autophagy

STRAP binds directly to JMY, and JMY promotes autophagosome formation, so we wondered whether STRAP also regulates autophagy. To test this idea we first over-expressed the STRAP protein in U2OS cells stably expressing GFP-LC3. We found that increased STRAP expression significantly reduces the average migration speed of LC3 vesicles (Figure 2G). We next quantified the degree of colocalization between JMY and LC3 in cells from which we had knocked out STRAP expression using CRISPR (Figure S4). Loss of STRAP significantly increased the degree of co-localization between JMY and LC3 in both fed and starved cells (Figure 3A-B), suggesting that STRAP negatively regulates autophagosome movement, possibly by competing with LC3 for binding to JMY.

**Figure 3.**
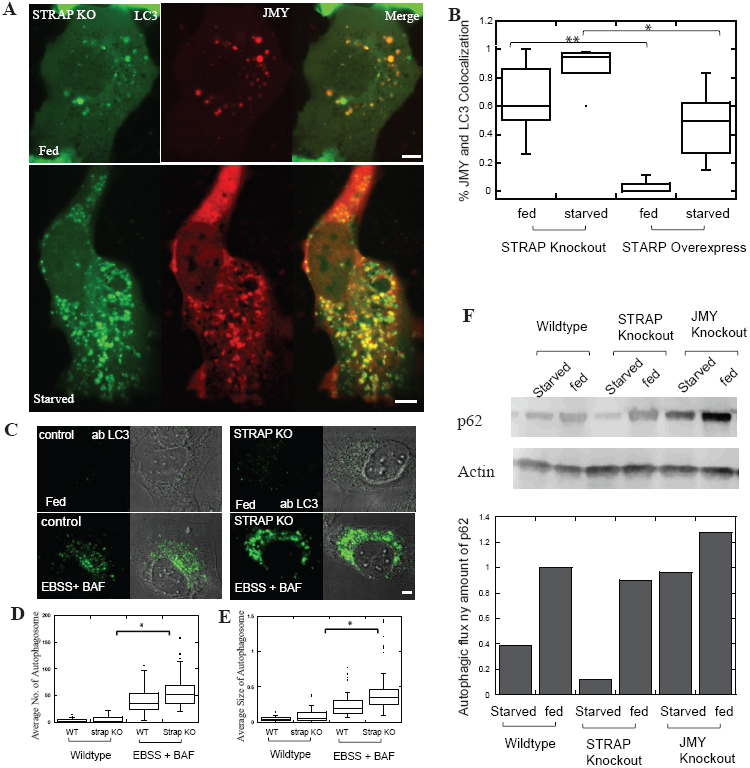
STRAP is a negative regulator of autophagy. (A) More JMY colocalizes with LC3 vesicles in STRAP knockout cell lines under both fed and starved conditions (B) Quantification of colocalization of JMY and LC3 in cells either lacking STRAP or over-expressing STRAP (n= 3 independent experiments, 15 ∼ 24 cells, **p < 0.01, *p < 0.05, student t test). (C) Immunofluorescent staining of LC3 in wildtype (WT) and STRAP knockout cell lines. U2OS cells were either fed (top) or starved in EBSS with 100 nM Balfilomycin A to block autophagosome maturation (bottom). (D-E) Autophagosome number and size increase significantly in STRAP knockout cells (n= 3 independent experiments, 60 ∼ 68 cells, *p < 0.05m, student t test). (F) Autophagic flux is increased in STRAP knockout cell lines, while it is decreased in JMY knockout cell lines. The autophagic flux is monitored by relative p62 protein concentrations measured by western blot. Temperature for live cell imaging: 37⁰. Scale bar: whole cell, 5 μm.

We next used our knockout cells to test whether STRAP functions as a negative regulator of autophagy. We used immunofluorescence to quantify LC3-positive autophagosomes in wildtype and STRAP knockout cells, and found that loss of STRAP expression significantly increased the number and size of endogenous autophagosomes under both fed and starved conditions (Figure 3C-E). We next tested whether loss of STRAP or JMY perturbs autophagic flux by quantifying cellular levels of p62, a cargo receptor and high-volume substrate for autophagy. Increased levels of p62 are associated with decreased autophagic flux and vise versa. In STRAP knockout cells we detected decreased amounts of p62, consistent with an increase in autophagic flux. Conversely, JMY knockout cells contained higher steady-state concentrations of p62 compared to wildtype cells, indicating a decrease in autophagic flux (Figure 3F). Together our results argue strongly that STRAP indeed functions as a negative regulator of autophagy.

### *In vitro* STRAP inhibits JMY’s intrinsic nucleation activity and its ability to activate the Arp2/3 complex

STRAP over-expression decreased motility of LC3-positive vesicles, so we used purified proteins to test whether STRAP directly regulates actin assembly by JMY. Using Forster Resonance Energy Transfer (FRET) to detect interaction between donor‐ and acceptor-labeled proteins, we found that purified STRAP binds full-length JMY with sub-micromolar affinity (K_d_ = 268 nM) (Figure 4A-B). Remarkably STRAP inhibits both the intrinsic actin nucleation activity of JMY (Figure 4C) and its ability to promote branched actin nucleation by the Arp2/3 complex (Figure 4D). In both sets of experiments inhibition by STRAP was concentration-dependent, with half-maximal effective concentrations (EC50’s) approximately equal to the measured equilibrium dissociation constant (K_d_). Although STRAP’s inhibition of JMY was incomplete ‐‐plateauing at approximately 80%‐‐ these experiments establish STRAP as the first known negative regulator of JMY’s actin assembly-promoting activities.

**Figure 4.**
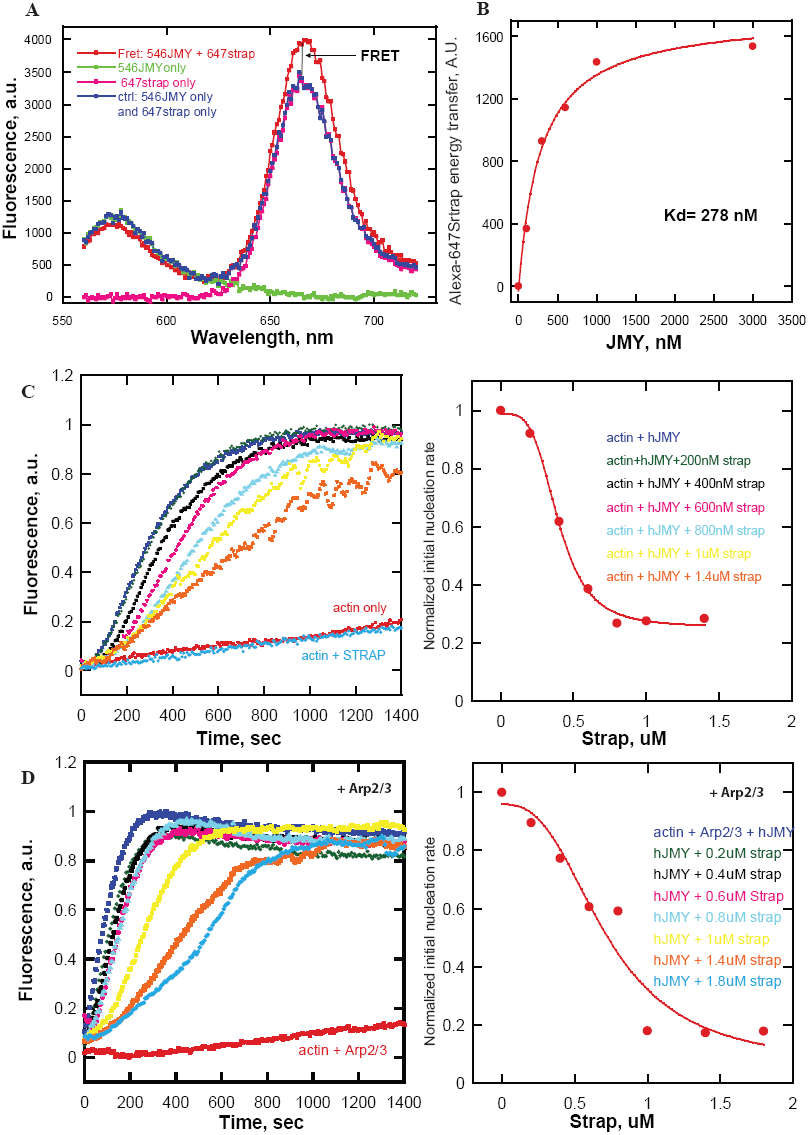
In vitro purified STRAP binds JMY and inhibits both intrinsic nucleation activity and the ability to activate the Arp2/3 complex. (A) Forster Resonance Energy Transfer (FRET) assay for the binding of purified STRAP to JMY. The curves are fluorescence intensity as a function of emission wavelength of 160 nM Alexa-546-labeled JMY (green) as donor, 600 nM Alexa-647-labeled STRAP (pink) as acceptor, and mixture of both fluorescently labeled proteins (red). The traces of donor alone and acceptor alone are combined to serve as baseline (blue) indicating no interaction. FRET is detected as an increase in acceptor emission intensity compared to baseline. (B) Binding isotherm for STRAP-JMY interaction derived from FRET assay (Kd=∼300 nM). Alexa-546-labeled JMY is titrated in 300 nM Alexa-647-labeled STRAP in 20 mM Hepes buffer (pH 7.4), 100 mM KCl. (C) STRAP inhibits JMY’s intrinsic nucleation activity. Actin assembly was monitored by pyrene-actin fluorescence in various concentrations of JMY and STRAP (left). The initial slope (first 250 seconds) of each curve was normalized and plotted as a proxy for nucleation rate (right). (D) Dose dependence of Strap inhibition of Arp2/3 complex activation by JMY. Pyrene-actin polymerization assay was performed in in 50 mM KCl, 1 mM MgCl_2_, 1 mM EGTA, and 10 mM Imidazole (pH 7.0) buffer with 1μM (5% labeled) actin, 200 nM JMY, 25 nM Arp2/3 as noted and STRAP as indicated.

### LC3 binds JMY, enhances intrinsic actin nucleation activity, and promotes activation of the Arp2/3 complex

Coutts et al. reported that mutation of the LC3-interacting region results in loss of JMY-associated actin structures in the cytoplasm (Coutts and La Thangue 2015). We also observed that vesicles containing both JMY and LC3 move through the cytoplasm on dynamic actin comet tails, so we hypothesized that LC3 might recruit JMY to membranes and stimulate its ability to make actin filaments. To test this idea, we first used a FRET-based binding assay to quantify the affinity of LC3 for full-length JMY and several JMY truncation mutants. We observed the highest affinity binding (K_d_∼55 nM) between LC3 and an N-terminal fragment of JMY spanning amino acids 1-314 (Figure 5A-B).

**Figure 5.**
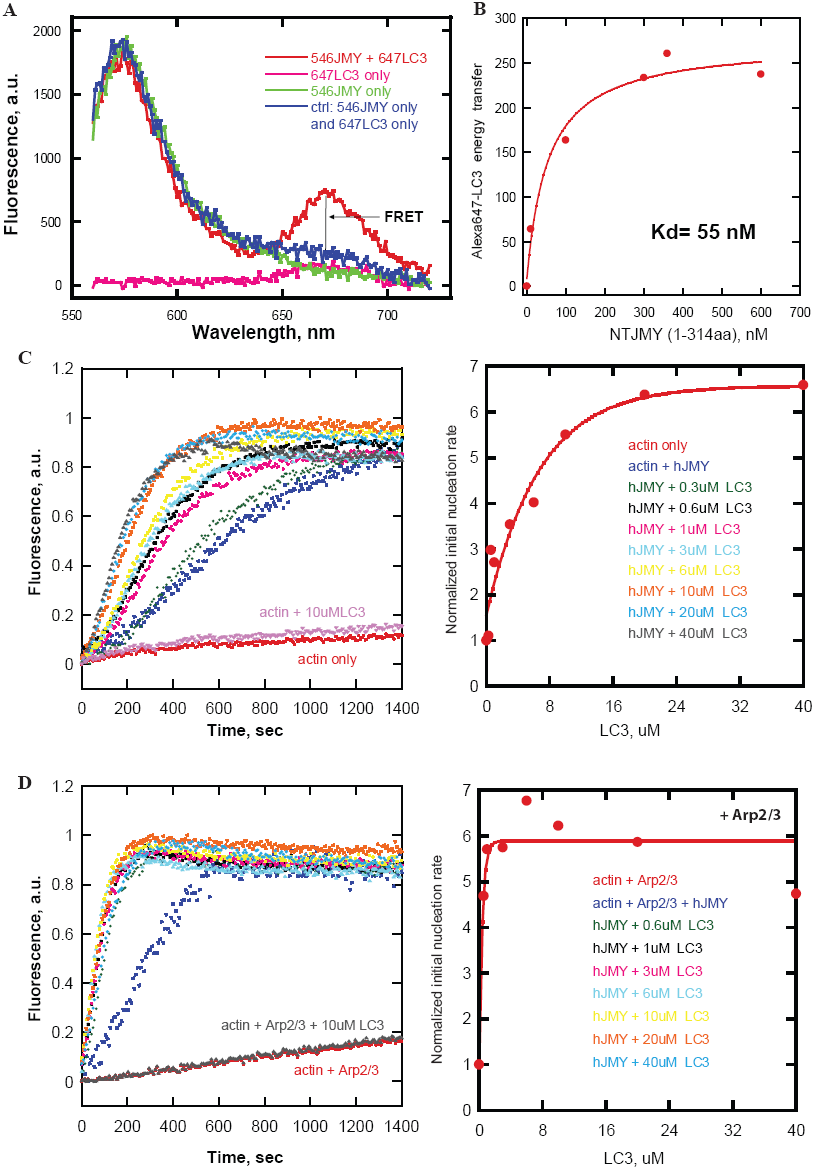
In vitro LC3 enhances intrinsic nucleation and Arp2/3 activation by JMY. (A) Forster Resonance Energy Transfer (FRET) assay for LC3 binding to JMY. The curves are fluorescence intensity as a function of emission wavelength of Alexa-546-labeled JMY (green) as donor, Alexa-647-labeled LC3 (pink) as acceptor, and mixture of both fluorescently labeled proteins (red). (B) Binding isotherm derived from FRET assay for the interaction of LC3 with the N-terminal region (residues 1-314) of JMY (Kd=∼50 nM). We added various concentrations of Alexa-546-labeled NT JMY to 200 nM Alexa-647-labeled LC3 in 100 mM KCl with 20 mM Hepes buffer (pH 7.4). (C) Dose dependence of LC3 stimulation of JMY’s intrinsic nucleation activity. LC3 promotes JMY’s intrinsic nucleation activity in pyrene-actin polymerization assay (left). The initial slope (first 250 seconds) of each curve is normalized and plotted as a proxy for nucleation rate (right). (D) Dose dependence of LC3 enhancement of Arp2/3 complex activation by JMY. Pyrene-actin polymerization assays were performed in 50 mM KCl, 1 mM MgCl_2_, 1 mM EGTA, and 10 mM Imidazole (pH 7.0) with 1 μM actin, 200 nM JMY, 25 nM Arp2/3 as noted, 16 mM NaCl, and LC3 as indicated.

We next tested the effect of LC3 on JMY’s intrinsic and Arp2/3-dependent actin nucleation activities using pyrene-labeled actin. As judged by the time-dependent changes in pyrene fluorescence, soluble LC3 had no effect on actin assembly in the absence or presence of the Arp2/3 complex. In the presence of full-length JMY, however, LC3 enhanced the intrinsic actin nucleation activity in a concentration dependent manner (Figure 5C). Similarly, LC3 accelerated actin assembly in the presence of both JMY and the Arp2/3 complex (Figure 5D).

To test whether membrane-associated LC3 also enhances JMY’s nucleation activity, we made liposomes with a combination of phosphatidyl choline (PC) and phosphatidyl serine (PS), doped with 2% NiNTA-conjugated 1,2-dioleoyl-*sn*-glycero-3-succinyl (DGS). We included Ni-conjugated lipids to recruit His-tagged LC3 and made liposome-bound LC3. This liposome-bound LC3 stimulates JMY’s intrinsic and Arp2/3-dependent nucleation activities in pyrene-actin assembly assays even more robustly (Figure 6A-B) than soluble LC3.

**Figure 6.**
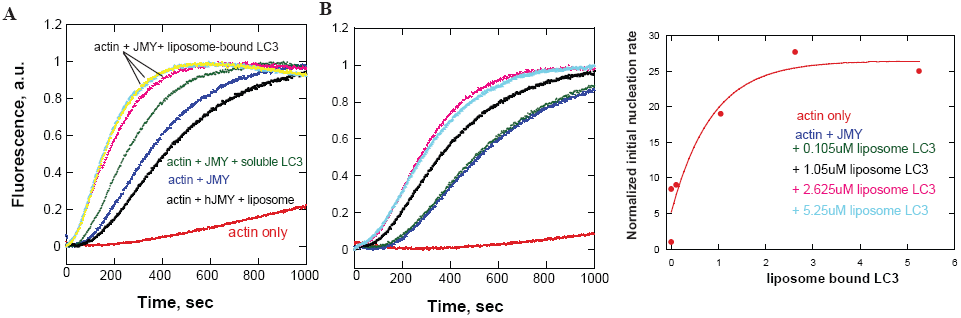
Liposome-bound LC3 stimulates JMY’s nucleation activity. (A) Liposome-bound LC3 promotes JMY’s intrinsic nucleation activity more potently than an equal amount of soluble JMY in a pyrene-actin polymerization assay. Time courses of actin assembly with liposome-bound LC3 (5.25μM) are shown in three replica (yellow, cyan, pink) to confirm no error in pipetting of liposomes. (B) Dose-dependence of liposome-bound LC3 enhancement of JMY’s nucleation activity (middle). The initial slope (first 250 seconds) of each curve is normalized and plotted as a proxy for nucleation rate (right). Pyrene-actin polymerization assays were performed at 23⁰ in 50 mM KCl, 1 mM MgCl_2_, 1 mM EGTA, and 10 mM Imidazole (pH 7.0) with 1 μM actin, 200 nM JMY, 5.25 μM soluble LC3, and liposome-bound LC3 as indicated.

### LC3 affects actin assembly via a cryptic regulatory sequence in the N-terminal region of JMY

We considered three possible mechanisms by which LC3 could enhance JMY’s ability to make actin filaments in both the absence and presence of the Apr2/3 complex. Firstly, we investigated a role for LC3 oligomerization, in part because previous work demonstrated that dimerization of Spire-family nucleators (Quinlan et al., 2007) and WASP-family nucleation promoting factors (Padrick et al., 2011) significantly increases their activity. When we measured the molecular weight of purified LC3 in solution using sedimentation equilibrium ultracentrifugation, however, we found that, similar to a related protein, GABARAP (Coyle et al., 2002), LC3 forms dimers only at high concentration (> 80uM). Given that the concentrations of LC3 used in our in vitro actin assembly experiments are well below the apparent K_d_ for dimer formation, LC3 oligomerization is unlikely to explain the enhancement of JMY activity. Secondly, we wondered whether JMY has a cryptic LC3 interacting region (LIR) buried somewhere within its actin regulatory sequence (PWWWCA). This hypothesis is appealing because intra‐ and inter-molecular contacts with Arp2/3-activating VCA sequences regulate the activity of other nucleation promoting factors (e.g. WASP and WAVE). In a FRET-based binding assay LC3 does not interact with the C-terminal fragment of JMY (CCPWWWCA, amino acids 315-983) that contains the Coiled-Coil, Proline-rich, and WWWCA domains (Figure S6B). In addition, we found that LC3 has no effect on actin nucleation or Arp2/3 activation by C-terminal JMY constructs (Figure S6A). These results fit with the previous identification of a single LC3-Interacting Region (LIR) near JMY’s N-terminus (Coutts and La Thangue, 2015).

Finally, we asked whether LC3 regulates nucleation activity via interaction with JMY’s N-terminal LIR. We began by testing whether JMY’s activities are inhibited by an intramolecular interaction, similar to those that regulate some formins (Li and Higgs, 2003) and WASP-family nucleation promoting factors (Rohatgi et al., 1999). Briefly, we tested whether truncation mutants containing the N-terminal region of JMY (NT-JMY, amino acids 1-314) can inhibit actin assembly stimulated by the C-terminal region (CCPWWWCA, amino acids 315-983). To our surprise, addition of the NT-JMY truncation mutant actually increases the rate of actin nucleation by the CCPWWWCA construct (Figure S6D). This increased activity turns out to reflect the contribution of a cryptic actin-nucleating region within the NT-JMY construct. Although this region contains no identifiable actin-binding motifs, we find that the NT-JMY truncation mutant by itself accelerates actin assembly in a concentration-dependent manner (Figure 7A-B). Moreover, addition of LC3 further enhances actin nucleation by NT-JMY (Figure 7C). This previously unsuspected nucleation activity suggests that NT-JMY contains one or more cryptic actin binding sites, which we confirmed by measuring fluorescence resonance energy transfer between NT-JMY and fluorescent actin monomers bound to the polymerization inhibitor Latrunculin-B (Figure S6E). We conclude, therefore, that LC3 promotes JMY-dependent actin nucleation by interacting with, and enhancing the activity of, a previously unknown actin regulatory sequence in the N-terminal region.

**Figure 7.**
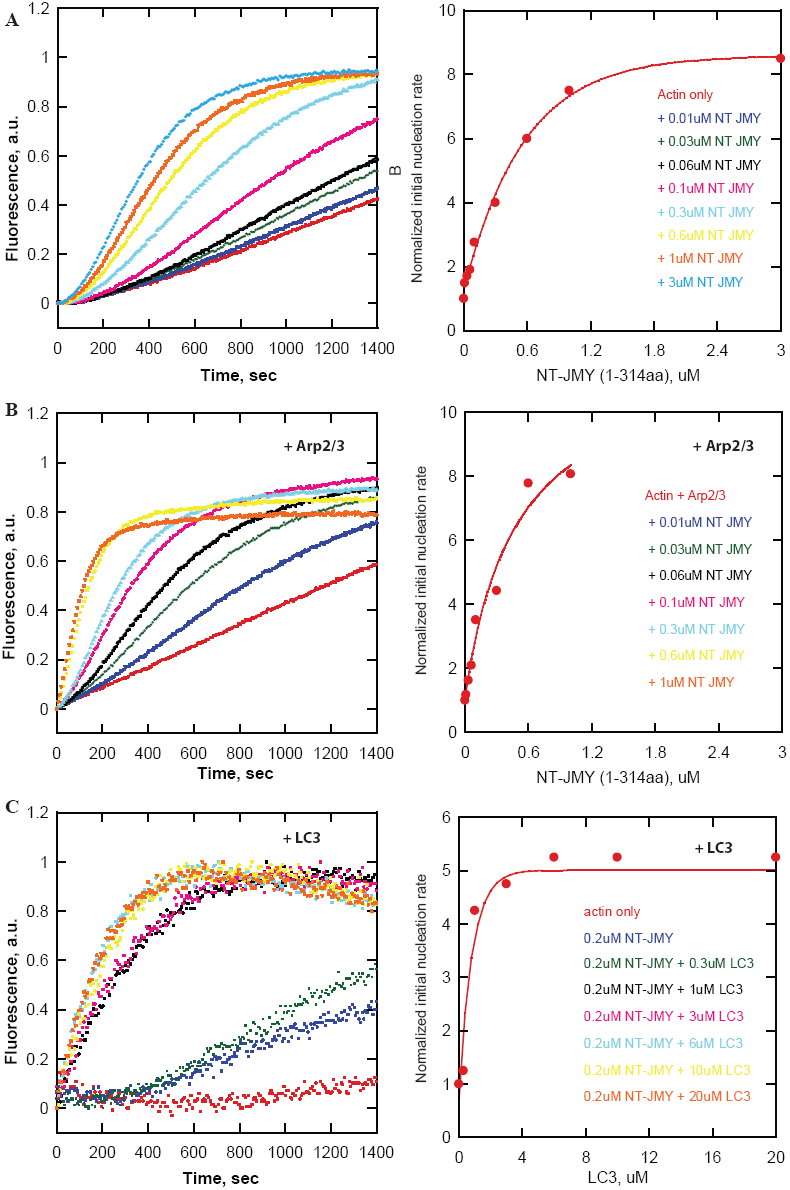
The effect of LC3 on actin nucleation and Arp2/3 activation by JMY reveals a cryptic actin regulatory sequence in the N-terminal region of JMY. (A-B) NT JMY (residues 1-314) promotes concentration-dependent actin nucleation in the absence (A) and presence (B) of the Arp2/3 complex. Actin assembly was monitored by the fluorescence of pyrene-labeled actin (left). The initial slope (first 250 seconds) of each curve is normalized and plotted as a proxy for nucleation rate (right). Pyrene-actin polymerization assays are performed at 23⁰ in 50 mM KCl, 1 mM MgCl_2_, 1 mM EGTA, and 10 mM Imidazole (pH 7.0) with 2 μM actin, 25 nM Arp2/3 and NT JMY as noted. (C) LC3 enhances NT JMY (residues 1-314) intrinsic actin nucleation activity. Actin assembly in the presence of various concentrations of JMY and LC3 was monitored by the fluorescence of pyrene-labeled actin (left). The initial slope (first 250 seconds) of each curve is normalized and plotted as a proxy for nucleation rate (right). Buffer conditions: same as in (A) with 1 μM actin, 200 nM NT JMY, 16 mM NaCl and LC3 as indicated.

### In vitro reconstitution of LC3‐ and JMY-dependent actin comet tail formation from purified components

Lastly, we asked whether LC3 is sufficient to recruit JMY to autophagosome membranes and stimulate actin-based motility by reconstituting the process *in vitro*, using purified components. We coated 4.5 μm glass microspheres with a combination of phosphatidyl choline (PC) and phosphatidyl serine (PS), doped with 2% NiNTA-conjugated 1,2-dioleoyl-*sn*-glycero-3-succinyl (DGS). We included Ni-conjugated lipids to recruit His-tagged LC3 that was labeled with a fluorescent dye, Alexa 647 (Figure 8A). By themselves, NiNTA-doped, lipid-coated microspheres did not recruit full-length JMY, labeled with Alexa 546 (Figure 8B). The LC3/lipid-coated beads did, however, recruit full length JMY and ‐‐in the presence of actin, capping protein, profilin, and the Arp2/3 complex‐‐ initiated the assembly of polarized, branched actin networks (Figure 8C) similar to the motile actin comet tails we observed in live cells. Interestingly, addition of soluble STRAP to this reaction does not prevent recruitment of JMY to the LC3-coated microspheres, but strongly suppresses formation a branched actin network (Figure 8D), consistent with STRAP’s ability to inhibit JMY’s actin-nucleating activity and with an inhibitory role for STRAP in JMY-mediated autophagosome movement.

**Figure 8.**
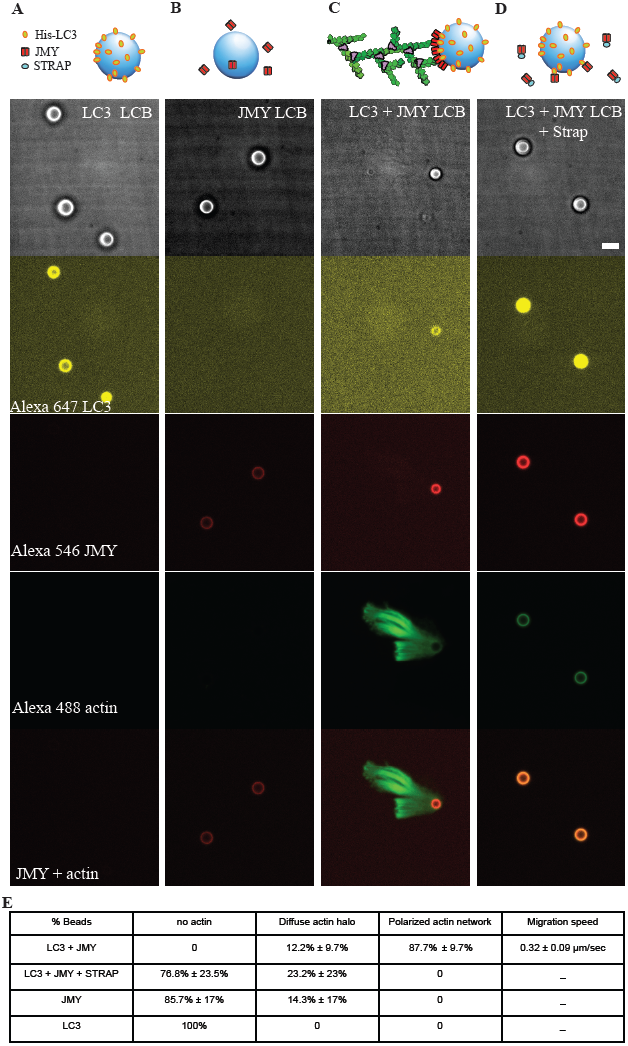
Reconstitution of LC3‐ and JMY-dependent actin comet tail formation from purified components. We mixed lipid-coated 4.5 μm glass micro-spheres with (A) his-tagged LC3; (B) full length JMY; (C) both his-tagged LC3 and JMY; and (D) STRAP together with his-tagged LC3 and JMY. From top to bottom: bright field images; Alexa-647-labeled LC3; Alexa-546-labeled JMY; Alexa-488-labeled actin; merger of fluorescent signal from Alexa-546-labeled JMY and Alexa-488-labeled actin. (E) Quantification of actin network on lipid-coated bead surface. Buffer conditions: 8 μM actin (10% labeled with Alexa 488), 200 nM Arp2/3, 400 nM Capping protein, 8 μM profilin, 2 μM STRAP as noted, 1 mg/ml BSA, 1 mg/ml β-casein, 50 mM KCl, 1 mM MgCl_2_, 1 mM EGTA, 0.2 mM ATP, and 20 mM Hepes (pH 7.0). Temperature: 23⁰. Scale bar, 5 μm.

## Discussion

Although it promotes actin filament assembly by multiple mechanisms (Zuchero et al., 2009), JMY was first described as a co-activator of p53-mediated apoptosis that accumulates in the nucleus in response to DNA damage. Early work identified several binding partners, including p300 and STRAP, which collaborate with JMY to promote apoptosis, but the regulation of JMY-mediated actin assembly has remained mysterious. Here we find that JMY’s nuclear partner, STRAP, also modulates its ability to create actin filaments in the cytoplasm. Previous work showed that DNA damage induces ATM-dependent phosphorylation of STRAP, leading to its nuclear localization and the formation of a complex between STRAP, JMY, and p300 (Demonacos et al., 2001). This is believed to be part of the reason JMY accumulates in the nucleus following DNA damage. We find, however, that in the absence of DNA damage STRAP also co-localizes with JMY in the cytoplasm on non-motile vesicles; and further that STRAP suppresses JMY’s actin nucleation activities *in vitro*. One consequence of this inhibition is that JMY likely cannot promote actin assemblywhen bound to STRAP inside the nucleus.

The best understood nucleation promoting factors are regulated by auto-inhibitory interactions within a single molecule (e.g. N-WASP) or a regulatory complex (e.g. the Wave Regulatory Complex). Upstream signaling molecules stimulate the activity of these NPFs by disruption auto-inhibition, either allosterically (Rohatgi et al., 2000; Padrick et al., 2008) or competitively (Okrut et al., 2015). The ability of LC3 to promote actin nucleation by JMY, however, does not involve disrupting an autoinhibitory interaction, but rather the enhancement of a previously unsuspected nucleation activity associated with JMY’s N-terminal LIR sequence. Although surprising, this result explains previous report that the N terminal region of JMY is essential for its nucleation activity in the cytoplasm (Coutts and La Thangue 2015).

We cannot rule out the existence of additional regulatory molecules, but based on our results we suggest a ‘two state’ or ‘two compartment’ model for JMY regulation. In this model, regulation consists of shuttling JMY molecules from a negative regulator on one membrane compartment to a positive regulator on a different set of membranes. Specifically, when cells are in the normal, ‘fed’ state JMY is sequestered and held in an inactive state by interacting with STRAP. During starvation-induced autophagy, LC3 titrates JMY away from STRAP and onto autophagosomal membranes, where its nucleation activity generates branched actin networks. Whether this shuttling involves formation of transient ternary complexes between STRAP, JMY, and LC3 or requires other proteins to mediate the transfer is a subject for further study.

Given that JMY and WHAMM both promote actin comet tail formation on autophagosomal membranes, why do mammalian cells require both nucleation promoting factors for efficient autophagy? A simple hypothesis is that the activities of WHAMM and JMY are required at different times and/or places. Autophagosome formation proceeds via multiple, morphologically and biochemically distinct steps: (1) localized initiation of a phagophore, (2) nucleation and (3) expansion of the phagophore membrane, (4) closure and (5) detachment of the autophagosome from the initiation site (Graef et al., 2013). Once detached, autophagosomes must move to and fuse with lysosomes/late endosomes. WHAMM is recruited to membranes by an early event in phagophore initiation: phosphorylation of phosphatidyl inositol in the endoplasmic reticulum to form PI(3)P. This acidic phospholipid binds and recruits WHAMM via interaction with PX domain-like sequences in the N-terminal region of the protein (Mathiowetz et al., 2017). In contrast, JMY interacts with LC3, whose lipidation happens after WHAMM is recruited to PI3P sites (Mathiowetz et al., 2017). This might suggest that JMY is strictly downstream of WHAMM, but the story is not so simple because we also observe colocalization and co-migration of JMY with proteins involved in the initiation and nucleation phases of phagophore assembly (DFCP1, ATG9) (Figure S5BC). These observations make it difficult to establish a clear difference in the timing of JMY and WHAMM recruitment during early stages of autophagosome formation. The fact that JMY, but not WHAMM, co-localizes with Lamp1 suggests that JMY activity might play a unique role in some late phases of autophagy.

The direct regulation of its actin nucleation activity by LC3 suggests that JMY plays a particularly important role in LC3-associated autophagic events, such as expansion of phagophore membranes; detachment of autophagosomes from sites of biogenesis; and fusion with lysosomes. This idea is consistent with the fact that JMY knockout mimics the effect of inhibiting the Arp2/3 complex, decreasing accumulation of autophagosomes in the perinuclear region when autophagic flux is blocked by Bafilomycin A (Figure S4D-F). This phenotype is consistent with failure to detach autophagosomes from parental membrane compartments and/or their inability to move and fuse with downstream compartments.

## Materials and Methods

### Constructs and reagents

Human JMY and mouse STRAP (also named as TTC5) derivatives were cloned into pHR vector with a C-terminal mGFP or mCherry or SNAP tag for mammalian expression. Human JMY full length and truncations were cloned into pFastBac with an N-terminal GST tag, followed by a prescission protease cleavage site and a SNAP tag for future labeling. Mouse STRAP full length was cloned into pet20b vector with a C-terminal 6 x His tag and an N-terminal KCK (Lysine-Cysteine-Lysine) tag for future labeling. Human EGFP-LC3B was purchased from addgene (11546) for mammalian expression, and human LC3 was cloned into pet20b vector with a C-terminal 6 x His tag and an N-terminal KCK (Lysine-Cysteine-Lysine) tag for future labeling. EGFP-DFCP1, GFPATG2, GFP-ATG9, GFP-VAPA were purchased from addgene. EGFP-F-Tractin, mTagBFP2-lifeact, and mCherry-Lamp1 were purchased from Davidson Lab Plasmid of UCSF imaging center.

### Cell lines, lentiviral infection and transfection

U2OS (ATCC) cells were cultured in DMEM media supplemented with 10% FBS, 2mM L-glutamine, and penicillin-streptomycin (Thermo Fisher). Lentiviral JMY-mGFP, JMY-mCherry and STRAP-mCherry were made from HEK293 cells, and infect U2OS cells. A U2OS cell line that stably expresses EGFP-LC3B was made by G418 screening and FACs sorting. Other mammalian expression constructs were transiently transfected into U2OS cells by using Lipofectamine 3000 (Invitrogen).

### CRISPR knockout and shRNA knockdown

The CRISPR knockout cell line of JMY is made by following the protocol described in Ran et al. (Ran et al., 2013). The knockout efficiency of JMY in single cell colony is validated by real time RT-PCR, western blot and immunofluorescence by using a homemade rabbit polyclonal antibody against hJMY. The sgRNA guide for JMY knockout is TCGCGCTCGTCGAACACATGGGG. The sgRNA used for STRAP knockout is ctttgacttgcatgctcaacagg. The knockout efficiency of STRAP is confirmed by real time RT-PCR (Biorad).

### Live-cell imaging

Microscopy was performed on an inverted microscope (Nikon Ti-E, Tokyo, Japan) equipped with a spinning-disk confocal system (Spectral Diskovery, Ontario, Canada), and imaged with a 60 x Apo TIRF Objective (NA 1.49) and EMCCDs camera (Andor iXon Ultra, Belfast, Ireland). For live unstarved cell studies, images were acquired at 37°C with 5% CO2 in Okolab stagetop incubator. Starved cells were imaged in HBSS (Sigma) supplemented with 20 mM HEPES and 100 nM Bafilomycin A. Images were captured at 1.0 s intervals at 16-bit resolution using Micro-manager software [Arthur Edelstein, Nenad Amodaj, Karl Hoover, Ron Vale, and Nico Stuurman (2010)]. Videos and images were prepared using the software Fiji (NIH). Migration speeds, and diameters were all determined using the manual-tracking, or manual measurement features of Fiji. To avoid observer bisa, we acquired and analyzed images double-blind.

### Protein expression, purification and labeling

Sf9 cells were cultured in insect media (Lonza Biowhittakar), and were infected by baculovirus of human GST-SNAP-JMY full length, GST-SNAP-JMY (1-314), and GSTSNAP-JMY (863-983). Insect cells were harvested at 72 hours after infection, and freshly lysed and purified. Using a microfluidizer, cells were lysed in 1x PBS, 1 mM EDTA, 10 mM β-mercaptoethanol, and 1 mM PMSF. The lysate were spun at 25K rpm (Beckman ultracentrifuge, Ti45 rotor) for 20 min, and supernatant were batch bound to glutathione sepharose 4b (GE healthcare). Resin was washed with lysis buffer containing 0.1% Triton X-100 and eluted with fresh made 33mM reduced glutathione in 50 mM Tris pH8.0. GST-SNAP-JMY was dialyzed and cleaved by GST-tagged prescission protease for 3 hours, and were rebound to glutathione sepharose 4b. The unbound fractions were collected and concentrated by dry sucrose and further purified by a gel filtration Superdex 200 column, followed by an anion exchange column MonoQ (GE healthcare). The pure untagged SNAP-JMY was dialyzed into storage buffer (20 mM Hepes, 100 mM KCl, 1mM EDTA, 0.5mM TCEP, 20% glycerol, pH7.4) and snap-frozen in liquid nitrogen. SNAP-JMY and derivatives were labeled with SNAP-cell-TMR-star (New England Lab) followed by the manufacturer’s protocol, and soluble free dye was removed with a G25 Sephadex column.

Expression of JMY-CCPWWWCA (315-983)-TEV-his6 construct was performed in BL21 Rosetta E. coli, induced with 30 μM IPTG at 18°C for 16 h. Using a microfluidizer, bacteria were lysed into 50 mM NaH2PO4, 300 mM NaCl, 10 mM imidazole, 10 mM β-mercaptoethanol, and 1 mM PMSF, pH 8. High speed supernatant was then batch bound to Ni-NTA resin (QIAGEN). Resin was washed with lysis buffer containing 20 mM imidazole and eluted with lysis buffer containing 200 mM imidazole. JMY-CCPWWWCA (315-983)-TEV-his6 was then dialyzed and cleaved by TEV protease for 3 hours, and was rebound to Ni-NTA. The unbound fractions were collected and further purified by a cation exchange MonoS column (GE Healthcare). Pure JMYCCPWWWCA were dialyzed into storage buffer (1 × PBS, 1mM EDTA, 0.5mM TCEP, 30% glycerol) and snap-frozen with liquid nitrogen before −80°C storage.

Expression of KCK-mSTRAP-his6 construct was performed in BL21 Rosetta E. coli, induced with 200 μM IPTG at 18°C for 16 h. Using a microfluidizer, bacteria were lysed into 50 mM NaH2PO4, 300 mM NaCl, 10 mM imidazole, 10 mM β-mercaptoethanol, and 1 mM PMSF, pH 8. High speed supernatant was then batch bound to Ni-NTA resin (QIAGEN). Resin was washed with lysis buffer containing 20 mM imidazole and eluted with lysis buffer containing 200 mM imidazole. mSTRAP was then dialyzed into 10 mM Hepes, pH 6.5, 1 mM EDTA, and 1 mM DTT. Proteins were then further purified by a cation exchange MonoS column (GE Healthcare). Pure mSTRAP were dialyzed into storage buffer (20 mM Hepes, 0.5mM TCEP, pH 7.4) and frozen with liquid nitrogen before −80°C storage.

Expression of hLC3-KCK-his6 was performed in BL21 Rosetta E. coli, induced with 50 μM IPTG at 18°C for 16 h. Using a microfluidizer, bacteria were lysed into 1 x PBS buffer, 10 mM β-mercaptoethanol, and 1 mM PMSF. High speed supernatant was then batch bound to Ni-NTA resin (QIAGEN). Resin was washed with lysis buffer containing 20 mM imidazole and eluted with lysis buffer containing 200 mM imidazole. hLC3 was then concentrated with 3KD Amico Ultra centrifugal filters (Millipore). Proteins were then further purified by a Superdex 200 gel filtration column (GE Healthcare). Pure mSTRAP were liquid nitrogen snap-frozen in storage buffer (20 mM Hepes, 50 mM NaCl, 0.5mM TCEP, pH 7.4) and stored in −80°C.

Labeling was achieved by combining reduced KCK-mSTRAP or LC3-KCK with 5 molar excess Alexa 546-maleimide (GE Healthcare) on ice for 15 min before quenching with 10 mM DTT. Soluble free dye was removed with a G25 Sephadex column. Labeling efficiency was assessed with a spectrophotometer.

Cytoplasmic actin was purified from rabbit skeleton muscle based on the methods described in Hu et al (Hu and Kuhn, 2012). Gel-filtered monomeric actin was stored in buffer containing 2 mM Tris, pH 8.0, 0.5 mM TCEP, 0.1 mM CaCl2, and 0.2 mM ATP. Actin was labeled on Cys-374 with Alexa Fluor 488 maleimide (Invitrogen) using the same method used for labeling KCK-mSTRAP. Arp2/3 complex is purified from Acanthamoeba castellani following the methods described (Dayel et al., 2001). Human profilin I was purified using established protocols (Kaiser et al., 1989). Recombinant mouse capping protein was purified using a protocol adapted from Palmgren et al. (Palmgren et al., 2001).

### Actin polymerization assays

Actin was labelled with pyrene iodoacetamide as described (Cooper et al., 1983), and stored on ice. For all assays, Arp2/3 was thawed daily and diluted with 1 mg/ml BSA in buffer A (0.2 mM ATP, 0.5 mM tris(2-carboxyethyl)phosphine (TCEP), 0.1 mM CaCl2, 0.02% w/v sodium azide, 2 mM Tris-HCl pH 8.0 at 4 °C). Actin polymerization assays were performed in 1 × KMEI (50 mM KCl, 1 mM MgCl2, 1 mM EGTA, 10 mM imidazole pH 7.0). Ca2+-actin was converted into Mg2+-actin by incubating actin in 50 mM MgCl2, 0.2 mM EGTA for 2 min before adding 10 × KMEI and test components. Pyrene fluorescence was measured with a Snergy4 platereader (BioTek). Unless otherwise noted, polymerization reactions contained 1 μM actin (labelled with 5% pyrene), 25 nM Arp2/3 and 200 nM JMY. To normalize fluorimetry data, we subtracted the offset from zero and then divided the plateau value of actin polymerization. For initial nucleation rate analysis, we fitted the first 250 sec of the kinetic data into linear plot, and normalize the slope by dividing the slope of actin only or actin plus Arp2/3.

### Fluorescence Resonance Energy Transfer (FRET)

We first scanned the donor and the acceptor separately, and the corresponding curves of donor and acceptor were added up to serve as baseline indicating no interaction. We then mixed the donor and acceptor, and FRET is indicated by the increase in acceptor emission intensity comparing with the baseline. Alexa 546-maleimide labeled mSTRAP or LC3 (donor) is titrated into 140 nM Alexa-647-SNAP labeled JMY (acceptor) in buffer with 20 mM Hepes, 100 mM NaCl, pH7.4. The FRET experiment is performed in Synergy 4 plate (BioTek) and ISS PCI/K2 fluorimeter.

### Liposome and lipid-coated beads preparation

To make lipid coated glass beads, glass beads (diameter = 4.5 μm, Bangs Technology) was first cleaned by 1 M HCl, 5 M NaOH, 1 mM EGTA, 70% ethanol, and pure ethanol in sequence and washed with MilliQ water between steps. The small unilamellar vesicles (SUV) was made by mixing 7.5 μmoles of PC (L-α-phosphatidylcholine), 2.3 μmoles of PS (L-α-phosphatidylserine), and 0.2 μmoles of Ni-NTA-DGS (18:1 DGS-NTA (Ni+2) 1,2-dioleoyl-sn-glycero-3-[(N-(5-amino-1-carboxypentyl)iminodiacetic acid)succinyl]), frozen and thaw for 20 times, and spin at 21,000 g for 30 min. I take the top 80% of supernatant and fill the tube with nitrogen gas. These SUV vesicles were further extruded through 100nm pore size filters to make liposome. Lipid-coated beads were made by mixing 7.5 μl of 10% bead slurry and 12.5 μl of 4μM SUV, bath-sonicated and rotated for 15 min, them washed 5 times with MilliQ water. The lipid-coated beads were stored in 20 mM Hepes pH 7.4, 150 mM NaCl for up to a week.

### Reconstitution of actin-based bead motility

To load proteins to lipid-coated beads, we mixed 4 μM 1% Alexa 647 labeled LC3-His6 and 4 μM 1% Alexa 546 labeled SNAP-JMY and rotate in cold room for 1hr. Then mix proteins with 20 μL lipid-coated beads in buffer including 1 mg/mL BSA, 1 mg/mL bcasein, 20 mM Hepes pH 7.0, 100 mM KCl and 0.5mM TCEP, and rotate for 10 min. Unbound proteins were removed by low speed centrifugation (200 g) for five times.

Labeled and unlabeled Ca-ATP actin were diluted to the desired labeled fraction, mixed 9:1 with 10 x magnesium exchange buffer (10 x ME: 10 mM ethylene glycol tetraacetic acid, EGTA, 1 mM MgCl2) and incubated on ice for 2 minutes to form 4 x final concentrations (4 × 8 μM) of Mg-ATP actin. We placed 8 μl of Mg-ATP actin at the bottom of a 1.5 ml Eppendorf tube and added 7 μl of motility protein mixtures (200 nM Arp2/3, 400 nM CP, and 8 μM profilin) and 1 μl of coated beads on the side of the tube. We washed both drops together with 16 μl 2 x buffer (2 x: 100 mM KCl, 2 mM MgCl2, 2 mM EGTA, 20 mM imidazole, pH 7.0, 200 mM DTT, 0.4 mM ATP, 30 mM glucose, 0.5% 1500 cP methylcellulose, 40 mg/ml catalase, 200 ug/ml glucose oxidase) and placed the reaction mixture in either a flow cell or slide-coverslip blocked in 1 mg/ml BSA.

#### Supplemental Materials

Figure S1 shows quantification of colocalization of JMY, LC3 and STRAP in cells expressing three proteins and a subset of JMY and STRAP positive puncta is enclosed by LC3 vesicle. Figure S2 shows LC3 and JMY-positive puncta colocalize on motile vesicles in U2OS cells expressing LC3 and JMY, while JMY and STRAP positive puncta are non-motile in cells expressing JMY and STRAP. Figure S3 shows JMY comigrate with ER-resident protein VAPA in an actin-propelled manner. Figure S4 shows test of JMY or STRAP knockout cell lines, total purified proteins used, and JMY-LC3 positive vesicles lost perinuclear enrichment when actin branched network is inhibited. Figure S5 shows wortmannin A treatment of cells and JMY colocalize with autophagy markers at different stages. Figure S6 shows biochemical analysis of JMY and related proteins.

Movie 1 shows JMY and STRAP colocalize on non-motile puncta in fed cell. Upon starvation, more JMY colocalize and comigrate with LC3. Movie 2 shows JMY‐ and LC3-positive puncta partially colocalize and comigrate. Movie 3 shows JMY‐ and STRAP-positive puncta partially colocalize and non-motile. Movie 4 shows polarized actin network propels JMY puncta movement. Movie 5 shows polarized actin network propels JMY and LC3 both positive puncta movement. Movie 6 shows CK666 inhibits JMY and LC3 both positive puncta movement. Movie 7 shows LC3 vesicles are much less motile in JMY knockout cell line.

## Acknowledgements

This work was supported by grants to RDM from the National Institutes of Health (R35-GM118119) and Howard Hughes Medical Institute. We thank members of the Mullins lab, especially Karen Cheng for CRISPR constructs, Sam Lord and Jiongyi Tan, for many constructive comments and helpful conversations. We also benefitted from advice and reagents provided by members of the Vale lab, especially Kara McKinley. RDM gratefully acknowledges the support and encouragement of Jenna Phillips and Carolina Mullins. XH thank Huadong Wang and Siqing Wang for their support and encouragement.

The authors declares no conflict of interest.

## Author Contributions

X. Hu and R.D. Mullins conceived the project; designed the experiments; and interpreted the results. X. Hu carried the experiments and data analysis. R.D. Mullins helped perform and analyze data from one experiment. Both authors discussed the results and wrote the final manuscript together.

